# Soil Microbial Composition and Structure Allow Assessment of Biological Product Effectiveness and Crop Yield Prediction

**DOI:** 10.1101/2021.02.09.430373

**Authors:** Nabeel Imam, Ignacio Belda, Adrian J. Duehl, James R. Doroghazi, Daniel E. Almonacid, Varghese P. Thomas, Alberto Acedo

## Abstract

Understanding the effectiveness and potential mechanism of action of agricultural biological products under different soil profiles and crops will allow more precise product recommendations based on local conditions and will ultimately result in increased crop yield. This study aimed to use bulk and rhizosphere soil’s microbial composition and structure to evaluate the effect of a *Bacillus amyloliquefaciens* strain QST713 inoculant on potatoes, and to explore its relationship with crop yield. We implemented NGS and bioinformatics approaches to assess the bacterial and fungal biodiversity in 185 soil samples, distributed over four different time points -from planting to harvest -from three different geographical regions in the United States.

In addition to variety, phenological stage of the potato plant and geography being important factors defining the microbiome composition and structure, the microbial inoculant applied as a treatment also had a significant effect. However, treatment preserved the native communities without causing a detectable long-lasting effect on the alpha- and beta-diversity patterns after harvest. Specific taxonomic groups, and most interestingly the structure of the fungal and bacterial communities (measured using co-occurrence and co-exclusion networks), changed after inoculation. Additionally, using information about the application of the microbial inoculant and considering microbiome composition and structure data we were able to train a Random Forest model to estimate if a bulk or rhizosphere soil sample came from a low or high yield block with relatively high accuracy, concluding that the structure of fungal communities is a better estimator of potato yield than the structure of bacterial communities.

**IMPORTANCE:** The manuscript’s results reinforce the notion that each crop variety on each location recruits a unique microbial community and that these communities are modulated by the vegetative growth stage of the plant. Moreover, inoculation of a *Bacillus amyloliquefaciens* strain QST713-based product on potatoes also changed specific taxonomic groups and, most interestingly, the structure of local fungal and bacterial networks in bulk and rhizosphere soil. The data obtained, coming from in-field assays performed in three different geographical locations, allowed training a predictive model to estimate the yield of a certain block, identifying microbiome variables -especially those related to microbial community structure- with a higher predictive power than the variety and geography of the block. The methods described here can be replicated to fit new models predicting yield in any other crop, and to evaluate the effect of any Ag-input product in the composition and structure of the soil microbiome.

## INTRODUCTION

Potato, the stem tuber vegetable produced by *Solanum tuberosum,* is the crop with the highest yield out of the five most important agricultural crops in the world (rice, wheat, soybeans, maize and potatoes). Although global production of potatoes in 2012 reached 364,808,768 MT, it has been calculated that actual yield corresponds to only about 10 to 75% of potential yield (1). Improving global agricultural crop production in a sustainable way is paramount given the current prospects for world population increase (2).

Potato yield has been directly correlated with edaphological and climate variation (3–5), with management practices (6) and with potato cultivar (7). Interestingly, the same biogeographical patterns have been identified as the main drivers of microbial community composition in potato plants (8–15), reinforcing the key role of soil microbiology in potato crop productivity (16).

Thus, in agro-ecosystems, the enhancement and sustainability of productivity can be assessed by means of the soil microbiome. Additionally, the Natural Resources Conservation Service of the US Department of Agriculture (17) links soil quality with the concept of soil health, acknowledging the relevance of soil microorganisms to drive soil functionality.

In this context, the use of substances, microorganisms, or mixtures thereof, known as plant biostimulants, is among the latest practices for sustainable food and energy production (18). Biological products are claimed to promote plant health and quality and recycling crop residues with low environmental impact (19, 20). Not surprisingly, the market for agricultural biological products is recording a CAGR of over 10% since 2017, and it is expected to reach a market size of over four billion dollars by 2025 (21). Rajabi-Hamedani and collaborators (22) argue that this growth is a consequence of the need to increase the efficiency of agrochemical inputs, to reduce crop damage caused by abiotic stress, and to reduce the environmental impact of production systems.

Most agricultural biological products based on microorganisms are expected to pertain to the functional group of Plant Growth Promoter species, so a direct impact in plant health (23, 24) and yield (25) is assumed. Different direct mechanisms involved in yield promotion have been demonstrated in certain bacterial strains, including: i) improving growth of tomato plants, by increasing root hairs development in a phytohormone-mediated process using an *Azospirillum brasilense* strain (26) or by increasing the tolerance to abiotic stresses through the action of an ACC deaminase produced by a *Burkholderia unamae* strain (27); ii) increasing plant growth by enhanced nutrient (P) acquisition in cucumber and tomato plants using a *Bacillus* sp. strain (28); iii) enhancing nodule formation by a two species consortia of *Pseudomonas bacteria survey performedputida* plus *Rhizobioum* sp. in beans (29); or by improving grain yield in rice by increasing panicle number through the use of an *Azospirillum amazonense* strain (30). In addition, some microbial strains have also shown an indirect effect in soil and plant health, as tools for *in situ* microbiome engineering, promoting the development of other beneficial microbial species, improving the resistance of the microbiome to the invasion of plant pathogens, and increasing the natural resistance of the plant against diseases (31).

Instead of assuming a simple, unidirectional and direct effect of a certain microbial strain in the physiology and development of plants, agricultural biological products face challenges with consistent field performance. Different strains and species can have different functional performance under specific environmental and ecological conditions (32). For this reason, biological products’ claims need to describe ecological and functional performance and not only be based on composition of matter (33).

In this work we aimed to contribute to global sustainability of the agricultural lands by demonstrating that assessment of bulk and rhizosphere soil microbial composition and structure can be practical tools to substantiate agricultural biological product claims, and at the same time they provide a toolkit for growers to assess and achieve increased yield and sustainability of their management practices. Applying “-omics” technologies we explored the subtle side effects of the microbial inoculant *Bacillus amyloliquefaciens* strain QST713, in the surrounding rhizosphere and bulk soil microbiota of potatoes, and its potential connection with the yield observed. We followed the recommendations of Ricci and collaborators (33) for field trials in one crop, in order to demonstrate that this product has *bona fide* effects. We were particularly interested in comparing the microbiome profile associated with treated vs. untreated samples over time and across diverse locations, to determine whether or not a common mechanism of action was at play. Both, the changes in the microorganism composition of samples across time as well as the evolution of the structure of the bacterial and fungal communities were assessed. Additionally, making use of potential correlations among microbiome profiles, product use and crop yield we built a yield prediction model as a first step towards guaranteeing growers the level of effectiveness of a product under different management and environmental conditions (weather, soil microbiome, soil type, crop variety, etc). Our observations conclude that individual microorganism abundances as well as the structure of the fungal and bacterial communities change slightly but significantly after application of the inoculant and that these changes can be associated with the unique yield response at each biogeographical location.

## RESULTS

In this work, we assessed bacterial and fungal communities of bulk and rhizosphere soil (soil health) of potato cultivars from three different regions of the United States (Sutton and Grant (Idaho), and White Pigeon (Michigan)). Our aim was to understand the effect of a microbial inoculant (*B. amyloliquefaciens* strain QST713) in the rhizosphere microbiota and its final legacy in the bulk soil microbiota after harvest. We were also trying to identify potential microbiome biomarkers associated with samples with low or high yields. A total of 185 samples from treated and untreated plots at each location were collected over four time points, from planting (T0) to harvest (T3), focusing on the early changes occurring after one (T1) and two (T2) months from planting, where T0 and T3 are bulk soil samples, and T1 and T2 are rhizosphere soil samples. Figure 1 shows that, in two of the three locations assayed the use of the inoculant had a significant effect on increasing the crop yield (Grant p-value 8.66×10^-10^, and Sutton p-value 7.67×10^-7^), without any detectable effect in the third location (White Pigeon p-value 0.31) which had, indeed, a much higher yield in both control and treatment samples.

**Figure 1.**
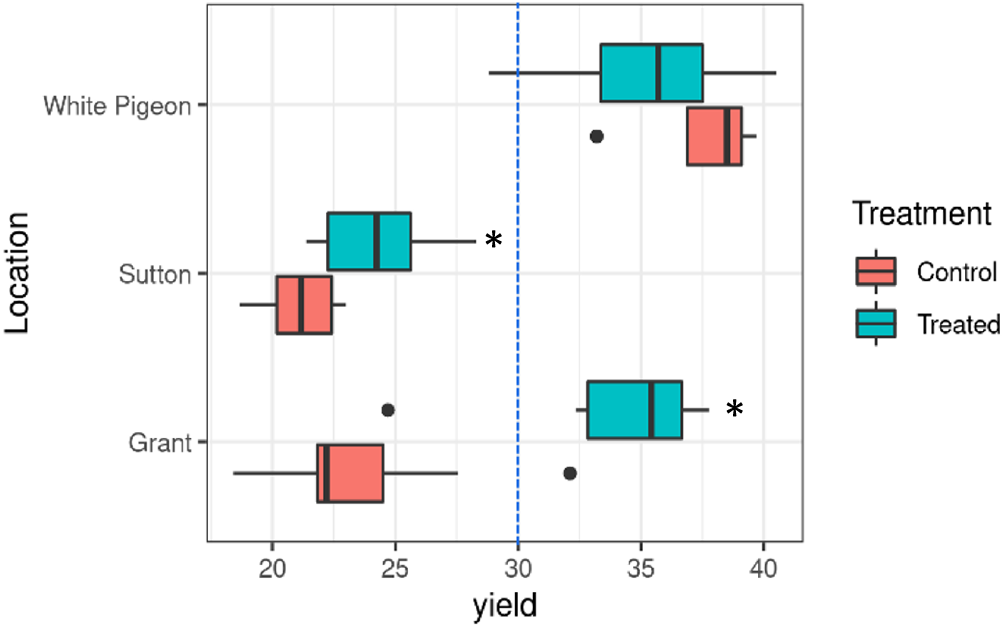
Yield data (t/ha) for control and treated blocks across locations. Discontinued line separates blocks into two categorical variables (≤ 30t/ha, > 30t/ha), and corresponds to one of the natural zero probability density points in the bimodal yield distribution. The box limits correspond to the 25th and 75th percentile, and the central line is the median. The whiskers are the 5th and 95th percentile. The dots represent outliers (points below 25th percentile - (1.5 * IQR) and above 75th percentile + (1.5 * IQR), where IQR is the interquartile range or absolute difference between 75th and 25th percentiles.

### Variety, phenological stage and geography drive the microbiome composition of bulk and rhizosphere soil of potato crops

Figure 2 shows a clear population dynamic occurring from T0 (before planting) to T1 and T2 samples (one and two months after planting, respectively) in all locations. Figure 2A shows that in terms of beta-diversity of bacterial populations, variety (R^2^=0.286), phenological stage (R^2^=0.286) and location (R^2^=0.042) had significant effects, with the treatment (R^2^=0.004) having a minor non-significant effect. However, for fungal populations (Figure 2C), variety dominates as the main driver of the beta-diversity patterns (R^2^=0.299), with phenological state having a much lower impact (R^2^=0.084) than in bacterial populations, and location (R^2^=0.067) and treatment (R^2^=0.007) having similar impacts to that in bacterial populations. Additionally, for fungal populations, all covariates had significant effects (full PERMANOVA data in Table S1). As shown in Figures 2A and 2C, White Pigeon is significantly different from Grant and Sutton; this can be easily explained by the geographical distance between locations, which correlates well with the Aitchison distances of samples in the PCoA analysis. There are also different edaphological and weather conditions at each of these locations, and a different crop variety in White Pigeon as compared to Sutton and Grant, all of which are major drivers of the soil microbial populations as previously observed by Rasche (10) and ínceoğlu (14) in potato soils. The significant differences between microbial community compositions before and after planting can be clearly seen at Figures 2A and 2C, where, despite the large differences between locations, T1 and T2 samples clustered in all the three locations, away from their respective T0, especially in the case of bacterial populations. Similar observations have been reported in maize (34), rice (35) and potato cultivars (13), and in forest soils (36).

**Figure 2.**
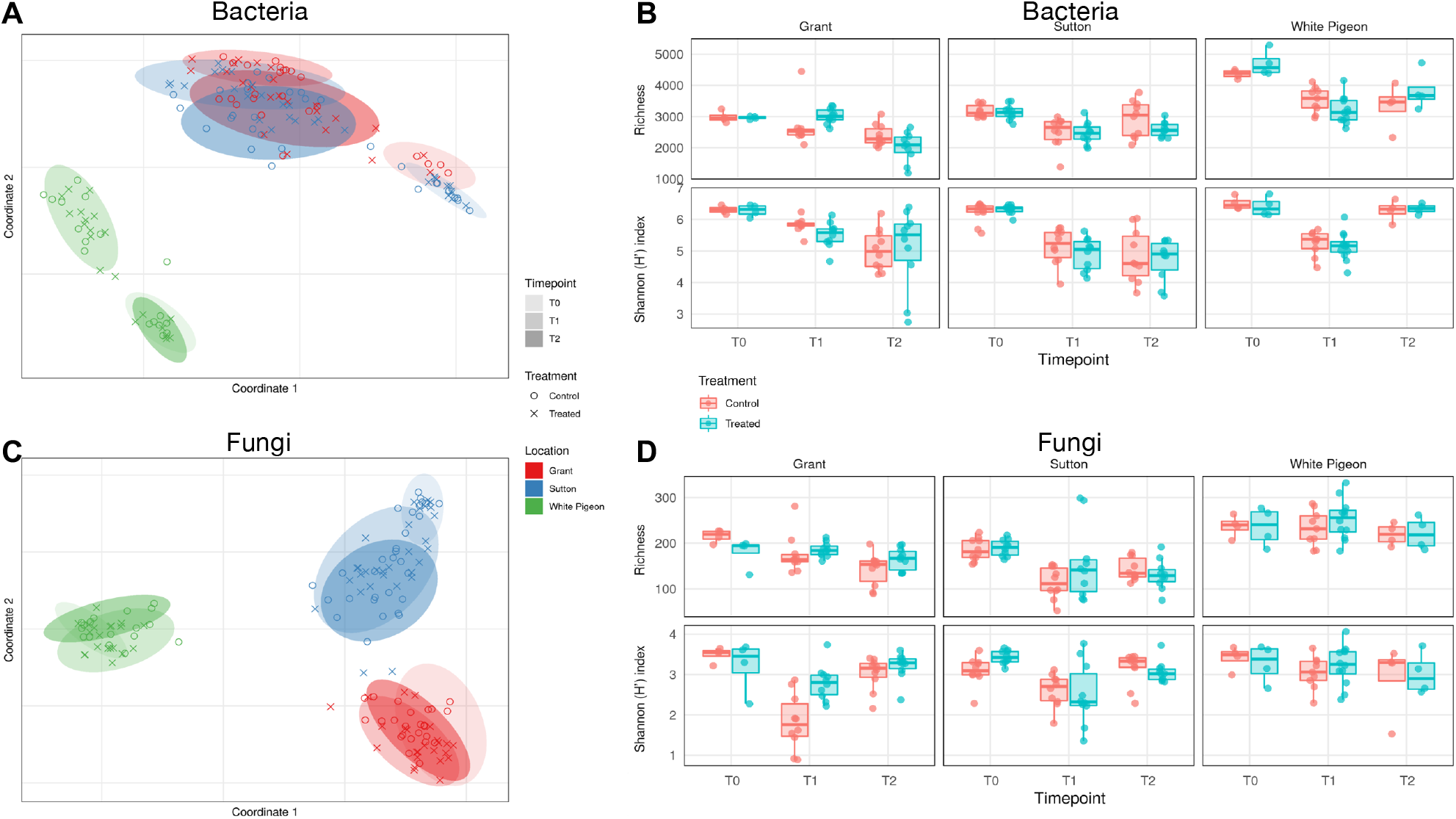
Beta- and alpha-diversity of bacterial and fungal populations in samples across locations and sampling times. **(A, C)** Beta-diversity (PCoA ordination) of bacterial and fungal populations. **(B, D)** Alpha-diversity (OTU Richness and Shannon (H’) index) of bacterial and fungal populations. T0 - before planting; T1 - one month after planting; T2 - two months after planting. Boxplot limits are the same as defined in Figure 1.

Regarding alpha-diversity (Figures 2B and 2D), there is a clear impact of planting in reducing the diversity of bacterial and fungal populations, as shown for both OTUs richness and Shannon (H’) index values from T0 to T1. This trend can be extended until time T2 in most cases -with the exception of the Shannon index for bacterial populations at White Pigeon and for fungal populations at Grant and Sutton-indicating that the phenological stage of the plant is one of the main drivers of changes at the alpha-diversity level in both bacterial and fungal populations. Additionally, comparing control versus treated samples at the same time point, we observed significant changes in Grant at T1 for bacterial richness and Shannon index as well as fungal Shannon index (Table S2). Interestingly, Grant was the site with the largest yield increase response due to treatment. When soil samples were again analyzed after harvest (T3) in Grant and Sutton locations, we observed that in spite of the marked microbial succession patterns found from T0 to T2, there was no significant changes in alpha-diversity between the microbial communities found in the soil before planting (T0) and after harvesting (T3) (Table S3); therefore, the plant’s associated soil microbiota seems to have cycled back to its original state. At the taxonomy level, despite clear population dynamic patterns from T0 to T2 sampling times in all the three locations and in both treated and untreated samples, samples from all three locations and times shared some of the most abundant genera for both bacterial and fungal communities (Figure S1). Figure S1 shows the top bacterial genera identified across samples in this study (core microbial species). Of these, five (*Arthrobacter, Pseudomonas, Sphingomonas, Streptomyces* and *Rhizobium)* also appeared in the soil bacteria survey performed by İnceoğlu (13) on potato fields. Among the top fungal genera shared across samples in our study (core fungal species) we found *Cryptococcus, Mortierella,* and *Alternaria.*

Thus, as previously reported (37) in tomato cultivars using a *B*. *subtilis* strain, and in soybean (38) and lettuce (39) cultivars using different strains of *B. amyloliquefaciens,* here we didn’t detect a durable impact of the treatment on the bulk soil microbial communities in terms of major taxa (Figure S1), and alpha- and beta-diversity (Figure 2), but instead we observed a clear temporal -cyclical-dynamics which differentiates bulk soil (T0 and T3) and rhizosphere soil (T1 and T2) samples (Figure 2).

### Elements of microbiome composition and structure can be effectively modulated by use of a *B. amyloliquefaciens*-based soil applied biological product

To dissect the specific effect of the biological product over the microbial composition across time at each location, we compared the fold change of each OTU in the treatment group from T0 to T1 (and from T0 to T2) *vs.* the fold change in the control group at the same time intervals per location (Table S4). Out of 17,241 unique bacterial OTUs in the samples of the study, 16 changed significantly from T0 to T1 (none in Grant, one in Sutton, and 15 in White Pigeon), and 100 from T0 to T2 (16 in Grant, 79 in Sutton, and five in White Pigeon). These OTUs belong to 73 genera, of which, 13 changed significantly in at least two locations: *Bacillus*, *Bradyrhizobium*, *Clostridium*, *Novosphingobium*, *Rhodoplanes, Sphingomonas, Sphingopyxis,* and *Woodsholea in* Grant and Sutton; *Agromyces, Flavobacterium, Pedobacter, and Sporosarcina* in Sutton and White Pigeon; and *Stenotrophomonas in* Grant and White Pigeon. For fungi, out of 1,702 unique OTUs, ten OTUs changed significantly from T0 to T1 (none in Grant, eight in Sutton and two in White Pigeon), and 32 from T0 to T2 (none in Grant, 32 in Sutton, and none in White Pigeon). These OTUs belong to 30 genera, of which, one changed significantly in at least two locations: *Cryptococcus* in Sutton and White Pigeon. Thus, despite variety, phenological stage and location having a larger effect than treatment in the composition of microorganism populations, the inoculant still generated common detectable abundance changes in at least two of the three locations for several taxonomic groups, some of which have known functionally relevant roles (*Bacillus, Bradyrhizobium*, *Flavobacterium*, *Pedobacter*, *Sphingomonas*, and *Stenotrophomonas*).

In order to get a deeper understanding of how the structure of the bacterial and fungal communities, and therefore the ecological relationships among microorganisms, impacts the effect of the bacterial inoculant, we studied the co-occurrence and co-exclusion patterns between pairs of OTUs in each sample of the trial. As some of us reported in a recent work (40), by studying the network properties of local communities inferred from the co-occurrences and co-exclusion patterns of a reference metacommunity it is possible to estimate ecological emergent properties (i.e. niche specialization, level of competition) of interest for the understanding of microbiome functioning. We first built metacommunities based on all samples of the trial. As an initial filter, for bacteria, we retained OTUs that were detected in at least 30% of the entire dataset, and 90% for fungal communities. This is due to the disproportionate number of unique OTUs detected in 16S *vs.* ITS soil sequencing. To keep the overall size of the data manageable we limited the number of selected OTUs to 4,000 with a maximum of 10 million possible significant pairs. We also filtered out OTU pairs that were not significantly (p < 0.05) enriched (co-occurrence) or depleted (co-exclusion). This resulted in metacommunity networks consisting of 3,339 nodes for bacteria (19.4% of the total 17,241 bacterial OTUs) and 447 nodes for fungi (26.3% of the total 1,702 fungal OTUs), which on average captured 92.11% of the bacterial abundance and 98.62% of the fungal abundance of the samples in the study. We then explored the structure of local microbiome communities, based on just the nodes present in each individual sample, aiming to detect changes in network properties that are associated with the application of the biological product at a specific location over time. Specifically, for the co-exclusion and co-occurrence bacterial networks, we calculated the modularity (a measure of the strength of partitioning of a network into modules) and transitivity (measure of the degree to which nodes in a network cluster together) as well as the proportion of co-exclusions and co-occurrences present in the local network compared to the total number of possible combinations among all OTUs in the sample.

Figures 3A and 3B show the evolution from T0 through T2 of four of the six local network properties studied across locations, for bacterial and fungal populations, respectively. Figure 3C lists those changes that have been significant (see Table S5 for full data) in time -from T0 to T1, and from T0 to T2-in treated vs. untreated blocks. In Grant there is a significant decrease in fungal co-occurrence transitivity and bacterial co-occurrence proportion from T0 to T1 in the treated samples when compared to untreated ones. In agreement with the observations of Ortiz-Alvarez (40) on their extensive survey of vineyard soils, it seems that any human intervention in a crop alters the structure of microbial communities of the soil, and a decreased transitivity on the fungal co-occurrence network seems to be a common indicator of these types of alterations. In the above-mentioned work, some of us argued that low clustered communities (those with low transitivity scores) can be associated with highly competitive environments with a high degree of niche specialization, which are among the most relevant properties of an ecosystem when trying to understand its functionality and its response to human interventions and land-use changes (41). It is also interesting to see a lagged effect (at T2) of the treatment in modifying some network properties of the bacterial communities in both Grant and Sutton. In Grant, the bacterial co-occurrence proportion increases from T0 to T2 (in contrast to the decrease from T0 to T1), and at the same time the transitivity of the bacterial co-occurrence network increases. In Sutton, both the bacterial co-occurrence proportion as well as the bacterial co-exclusion proportion increased from T0 to T2. Thus, when attending to the microbiome structure changes caused by the treatment in Grant and Sutton, which were the locations where treatment had a significant effect over yield, we can highlight significant treatment-mediated effects over the fungal and bacterial community networks that decreased from T0 to T1, and then increased in T2. Interestingly, and contrary to what was observed in Grant and Sutton, in White Pigeon, the location where treatment didn’t have a significant effect over yield and that had a different variety of potato, there was an increase in the bacterial co-exclusion modularity from T0 to T1.

**Figure 3.**
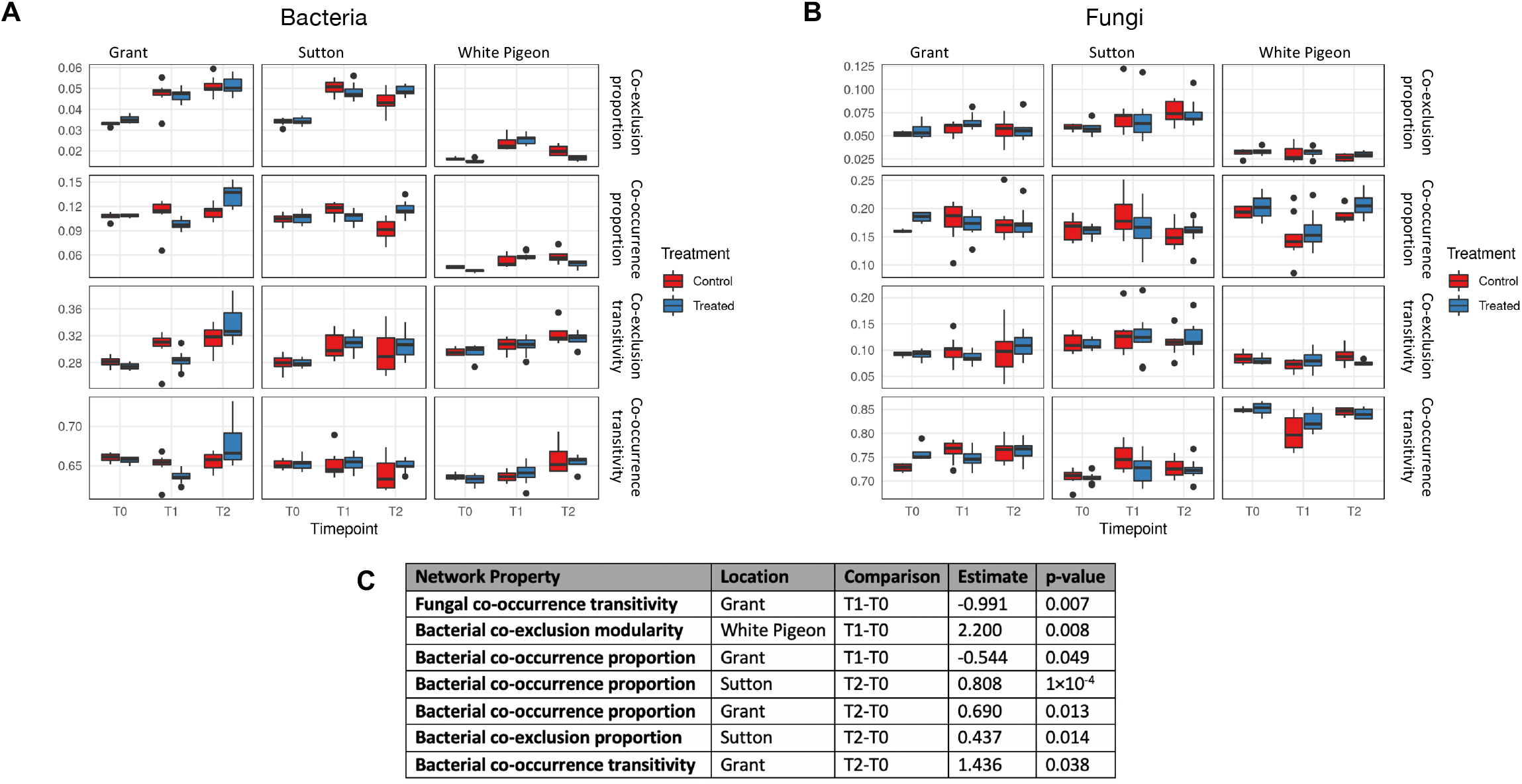
Local network properties across locations and sampling times. **(A, B)** Local network properties of bacterial and fungal populations in samples from the three locations (Grant, Sutton and White Pigeon) at three sampling times (T0 - before planting; T1 - one month after planting; T2 - two months after planting). **(C)** Significant changes from T0 to T1 and from T0 to T2 in treated vs. untreated blocks.

### Elements of microbiome composition and structure allow prediction of potato yield

We fitted a Random Forest model aiming to predict if a rhizosphere or bulk soil sample comes from a block with a yield ≤ 30t/ha or > 30t/ha, based on its microbiome composition and structure using multivariate compositional data (Principal Components from a beta-diversity ordination) and local network properties. We measured yield data in 20 treated and 20 untreated plots from the three geographical locations, and for each we utilized all samples available over times T0, T1 and T2. In total 112 samples were used for this task split into a training set of 84 samples and a test set of 28 samples. The result of this model showed a predictive accuracy of 78.6% (Figure 4A) and identified four variables (two network properties and two compositional) as the most important predictors of yield (Figure 4B), even with a higher importance than a variable we used to encompass the effects of geography and variety that are not accounted for by the microbial composition and structure. Surprisingly, the structure of fungal communities (i.e. fungal co-occurrence transitivity and co-exclusion proportion), showed a much higher predictive value than the structure of bacterial communities (Figure 4B, see Table S6 for full data on the importance of variables to the yield prediction model). We observed an inverse correlation between the co-occurrence transitivity of bulk and rhizosphere soil fungal communities and the yield found in the potato cultivars. This is a particularly important observation for understanding the effect of the *B. amyloliquefaciens-based* biological product assayed here in shaping the structure of fungal communities as a potential mechanism of action when increasing the yield. As shown in Figure 3B and Table S5, in going from T0 to T1 the increase in fungal co-occurrence transitivity in Grant is greater in the control samples than the treated ones, and this difference is significant (p-value=0.007). In Sutton -where a smaller but significant effect of the treatment increasing yield was also found (Figure 1)- in going from T0 to T1 there was a smaller increase in fungal co-occurrence network transitivity in the treated samples when compared to the control ones (albeit the difference is not significant, p-value=0.086). In White Pigeon instead, where the treatment did not have an effect over yield and where there was a different potato variety, there is a decrease in fungal co-occurrence network transitivity in going from T0 to T1 in treated samples, and even a more marked decrease in control samples.

**Figure 4.**
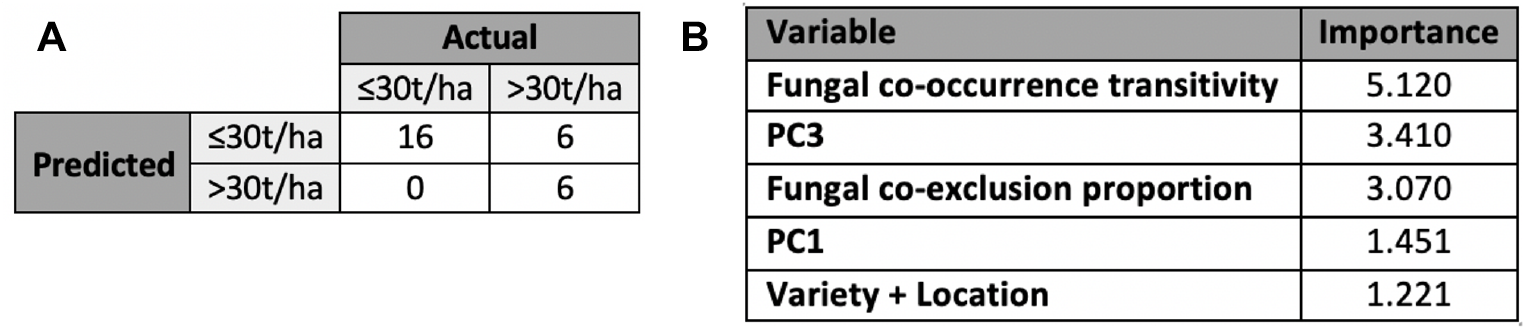
Random Forest yield model fitted to predict blocks with yields of ≤ 30t/ha or > 30t/ha based on soil microbiome composition and structure data. **(A)** Confusion matrix for the Random Forest model over the test set samples. **(B)** Importance figures of the main variables contributing to the predictive power of the Random Forest model.

The other two compositional variables (PC3 and PC1) contributing to the predictive power of the model fitted can be explored by looking at the taxonomy of the OTUs in each showing a significant correlation with the yield. It is necessary to keep in mind that the predictive power of PC3 and PC1 variables, as principal components of a multivariate analysis, came from the interaction patterns among the OTUs and not from their individual behavior. However, we can highlight the presence, for instance, of the fungal biocontrol agent *Trichoderma* sp. (42) as the OTU with the highest positive correlation with yield in PC3 (Figure S2).

As described in the methods section, since yield is constant for all samples within a plot, we converted yield to a categorical variable (≤ 30t/ha, > 30t/ha). The distribution of the yield data was bimodal, and thus it seemed logical to divide the categories on a zero probability density point for the bimodal distribution. However, in order to assess if this decision may have had an impact in the features identified as important by the yield predictive model presented here, we investigated the models resulting from splitting the yield data into three (≤ 26t/ha, > 26t/ha to ≤ 35t/ha, > 35t/ha) or four (≤ 20t/ha, >20t/ha to ≤ 26t/ha; > 26t/ha to ≤ 35t/ha, > 35t/ha) categories. As can be seen from Table S6, fungal co-occurrence transitivity and fungal co-exclusion proportion always had higher importance than geography and variety, independent of the number of yield categories used. In the model with three yield categories the bacterial co-exclusion proportion also had higher importance than geography and variety, whereas in the model with four yield categories, fungal co-inclusion modularity and PC12 had higher importance than geography and variety. However, dividing yield into more categories resulted in decreased accuracy (64.3% when splitting into three categories and 57.1% when splitting into four categories) due to the limited training set size being divided into an increasing number of categories.

Van Klompenburg and collaborators (43) performed a systematic literature review to identify the most used machine learning algorithms for crop yield prediction as well as the most used features to train those algorithms. They identified that most researchers have used neural networks in their work with the most frequently used features being temperature, rainfall and soil type. Interestingly, none of the articles reviewed utilized soil microbial or fungal composition or structure as features. In recent work, Jeanne and collaborators (16) developed a model to correlate potato yield to soil bacterial diversity. They showed that their species balance index related to potato yield (SBI-py) had a high correlation (0.77) with yield, whereas the Shannon diversity, Pielou diversity and Chao 1 diversity failed to correlate well with yield. Here, we built a machine learning potato yield model based on bacterial and fungal communities of rhizosphere and bulk soil and their structure, which can predict with relatively high accuracy whether a potato plot will have a yield of more or less than 30t/ha, which was the value that divided the bimodal distribution of yield in our training set. It is also worth noting that the dataset in this study included as a variable the application of a bioinoculant, thus this yield model also represents a first step towards understanding when and where biological products work. Despite the small sample size and the treatment of yield as a categorical value, independent of the number of categories used for splitting yield, we always found that the structure of fungal communities was a better estimator of potato yield than the structure of bacterial communities, which is a finding that merits further investigation.

## DISCUSSION

The use of microbial inoculants to increase the yield of plants is a useful strategy, increasingly used in agriculture. In addition to the direct impact of the microbial inoculant in the plant, due to its unique metabolic properties, the introduction of an allochthonous strain in the microbial rhizosphere and bulk soil ecosystems may have an impact on the entire microbiome, affecting the composition and structure of the native communities. Our work demonstrates that variety being the main driver of the microbial profile of rhizosphere and bulk soils from potatoes, phenological stage of the plant and geography also have a major impact in the microbiome composition, especially in the bacterial community. Even though relegated to last position, the use of a microbial inoculant based on *B. amyloliquefaciens* QST713 -a strain isolated from the soil of a Californian organic peach orchard with a demonstrated effective broad-spectrum bactericide and fungicide activity (44) through a number of different mechanisms of action (45, 46)- had a significant effect over the beta-diversity of fungal communities. Looking at alpha-diversity we observed significant changes at T1 in one location (Grant) for both bacterial and fungal communities in treated plots. Given that variety, plant phenological stage and geography have such strong influence over bulk and rhizosphere soil community composition and structure, the treatment effects observed were analyzed per location as evolution between two time points when comparing treated versus untreated plots. This also means that the patterns identified here as derived from product use may be of a more correlative than deterministic nature. Nonetheless, several OTUs changed significantly from T0 to T1 and from T0 to T2 in the inoculated soils, including several functionally important members of the soil microbiota, as well as modified specific microbial network properties. Specifically, a potential link between the bioinoculant and yield whereby the bioinoculant reduces the transitivity of the co-occurrence fungal network of the rhizosphere and bulk soil where it is applied through its biofungicide activity seems fit. Importantly, we also observed that this *B. amyloliquefaciens* QST713 trial did not cause any legacy effect on the microbiome profile of the soil analyzed after harvesting, i.e. the effect of the bioinoculant is cyclical and the native microbiome returns to its original state after harvesting. We also presented here a Random Forest yield prediction model for potatoes based on a soil health assessment of its microbial composition and structure. This model is our first step towards understanding not only why, but also when and where biological products work increasing yield. In addition, the significant contribution of the local network properties on the estimation of the actual yield of a certain block reinforces the idea of the need of a more functional vision of agriculture microbiomes, as certain emergent properties can be deduced from them. In particular, low clustered (low co-occurrence transitivity) fungal communities, as found here as positively contributing to the yield, are expected to be driven by a higher degree of niche specialization (40). Thus, *B. amyloliquefaciens* QST713 seems to help the soil microbiota adopt a conformation with lower fungal co-occurrence network transitivity than expected from untreated plots which is conducive to improved yield, but in a reversible manner (the fungal communities return to their original stage post-harvest).

Our model trained in only three locations, including only two potato varieties, and where half of the samples were treated with the bioinoculant may be biased, for instance, in recognizing the effects of *B. amyloliquefaciens* QST713 over the soil microbiome as the main features predictive of yield. However, the fact that fungal co-occurrence network transitivity is linked with potato yield, has not been reported before and merits further study. Possible future avenues of research derived from the current work include: i) investigating whether fungal co-occurrence network transitivity continues to be an important variable in models predicting yield in a more diverse datasets than the one described here (more varieties, more locations, more samples, wider variety of edaphological and weather conditions); ii) building predictive models containing not only microbiome data, but also edaphological and climate information; iii) investigating further the link between decreased fungal co-occurrence network transitivity and nutrient metabolism in the soil; and iv) investigating further metacommunity networks or individual sample networks with low transitivity to understand the taxonomic composition of modules and explain further each of the niches identified within them; among others. Ultimately, the inverse correlation between crop yield and fungal co-occurrence transitivity identified in this study as the potential mechanism of action of *B. amyloliquefaciens* QST713 in increasing potato yield is a useful concept to design and test interventions for increasing crop yield. This finding also demonstrates that assessment of soil microbial composition and structure in agricultural input trials can be practical tools to substantiate biological product claims, and that they provide a toolkit for growers to assess and achieve increased yield and sustainability of their management practices.

## MATERIALS AND METHODS

### Field trials

Russet Burbank potatoes were planted in Sutton and Grant locations in Idaho. Seed variety Lamoka was planted in White Pigeon, Michigan. Applications were performed at 46.770 l/ha spray volume combining grower standard practice of Quadris and Admire Pro plus the biological treatment, tank mixed and applied in-furrow. Treatment consisted of a biological product containing a minimum of 2.7×10^10^ CFU/g of *B. amyloliquefaciens* QST713 at a dose of 0.935 l/ha out of the total spray volume. Grower standard fungicide and insecticide applications were chemigated over trial area as seeded. Plots were 0.405 hectares (1 acre) each. Harvest was conducted by harvesting 2.787m^2^ within each plot. Yield weights were evaluated and recorded in lbs and cwt/ac.

### Sample collection

Whole plants from the field were collected and processed to obtain bulk soil and rhizosphere samples over the three regions. The field samples were processed to obtain bulk soil from all the root surfaces by vigorous shaking, and to collect rhizosphere samples we followed the protocol by Lundberg and collaborators (47) with slight modifications: roots (separated from mother tubers) were chopped into small bits and collected in a clean tube, filled with 40 ml of PBS buffer, vortexed and centrifuged to obtain a pellet. The rhizosphere pellet was stored at −80°C until genomic DNA was extracted. Samples were collected at four different time points: before planting and/or treatment (T0), one month after planting (T1), two months after planting (T2) and at harvest (T3). Samples from White Pigeon were collected only for three time points: T0, T1 and T2. From each time-point, a total of 20 samples (ten treated and ten untreated) were collected, except for White Pigeon at T0 (four treated and four untreated), T1 (12 treated and nine untreated), and T2 (four treated and four untreated), and for Grant at T0 (four treated and four untreated). A total of 76 bulk soil samples (T0 and T3) and 109 rhizosphere samples (T1 and T2) were collected. Samples were collected across different locations for each field and the composite was submitted for analysis, in order to achieve a more homogenized sampling reducing the effect of microbial variability.

### Sample analysis

After collection, samples were immediately sent for molecular analysis to Biome Makers laboratory in Sacramento, US. DNA extraction was performed with the DNeasy PowerLyzer PowerSoil Kit from Qiagen. To characterize both bacterial and fungal microbial communities associated with bulk soils and rhizosphere samples, the 16S rRNA and ITS marker regions were selected. Libraries were prepared following the two-step PCR Illumina protocol using custom primers amplifying the 16S rRNA V4 region and the ITS1 region described previously (48). Sequencing was conducted in an Illumina MiSeq instrument using pair-end sequencing (2×300bp). The bioinformatic processing of reads included the merging of forward and reverse paired reads to create robust amplicons, using Vsearch (49) with minimum overlaps of 100 nucleotides and merge read sizes between 70 and 400 nucleotides. OTU clustering was performed at 97% sequence identity, followed by quality filtering through *denovo* chimera removal using the UCHIME algorithm (50). Taxonomic annotation was performed using the SINTAX algorithm (51), which uses k-mer similarity to identify the top taxonomy candidate, after which we retained results where the species level had a score of at least 0.7 bootstrap confidence. We used the SILVA database version 132 (52) and UNITE database version 7.2 (53) as taxonomic references.

### Alpha- and beta-diversity analysis

Exploratory analyses of 16S and ITS OTU counts were conducted separately using the R package vegan (54). Alpha- and beta-diversity were analyzed using OTU counts. Alpha-diversity metrics (Shannon and richness) were calculated and plotted across all covariates available. Wilcoxon rank-sum tests were performed to compare control and treated samples within location-timepoint subgroups. For beta-diversity, Kruskal’s non-metric multidimensional scaling was performed in conjunction with Aitchison distances. Relative abundances for OTUs as well as annotations at various taxonomic levels (genera, families, etc.) were used in the analyses. Permutational multivariate analysis of variance was performed on the Aitchison distance matrix, using all possible combinations of the variety, location, timepoint and treatment variables.

### Differential abundance

For all subsequent analyses, the zero counts in the data were replaced. Valid values for replacement were calculated under a Bayesian paradigm, assuming a Dirichlet prior. Non-zero values were then adjusted to maintain the overall composition (55). Differential abundance determination was carried out using the R package edgeR (56). For each OTU, the fold change attributable to the treatment across different times (e.g. T0 to T1) was calculated. This was done by conducting a hypothesis test separately for each location, measuring the fold change of a given OTU in the treatment group (from T0 to T1) vs. the fold change in the control group (from T0 to T1), and then repeating the test but using times T0 and T2.

### Calculation of local network properties

Meta-community networks were built for 16S and ITS data separately using the methods described by Veech (57) and Ortiz-Álvarez (40). In a nutshell, we first built a metacommunity network of all samples: this was done by estimating the co-occurrence and co-exclusion that would occur solely by chance for all possible OTU pairs, given the data. We selected OTU pairs that occurred significantly more than expected by chance to create the co-occurrence networks. Similarly, those that occurred significantly fewer times than expected by chance constituted the co-exclusion network. Local networks (single sample-level) were calculated by subsetting the metacommunity network for OTU pairs detected in each sample and estimating a local network. The R package igraph was used to calculate network properties: modularity, transitivity and proportion of co-exclusions and co-occurrences in relation to the total number of combinations among all OTUs in a sample (58). An adequate description of the ecological meaning of the different network properties calculated in this work can be found in the review work of Proulx and collaborators (59). Network properties were compared using a linear model. Using the network property as outcome, hypothesis tests were performed to compare timepoint differences in treated vs. control samples (analogous to the approach used for investigating differential abundances).

### Yield model

Yield data was first explored using medians and inter-quartile ranges (IQRs). Wilcoxon rank sum tests were performed on these yield data. The OTU counts were transformed using the centered log-ratio (CLR) transformation. CLR-transformed 16S and ITS data were jointly projected onto 70 principal components. Yield was modelled as the outcome of these 70 principal components, along with fungal and bacterial network properties, treatment, soil type (to distinguish between bulk and rhizosphere soils), and a variable that encompasses variety and geography, using a probability forest as described by Malley (60). Since the yield is a constant variable for all time points within a plot, the yield was converted to a categorical variable (≤ 30t/ha, > 30t/ha). The threshold for this division, 30 tonnes, was set at a zero probability density point for the bimodal distribution of yield. We used a total of 112 samples (all T0 through T2 samples in the study for which we had yield data) and split them into training (n = 84) and test (n = 28) sets. Variable importance for each variable in the model was calculated using the Gini index. As a sensitivity test, probability forests were fit for a three-way split of the yield variable, and variable importances were compared. Among the 70 principal components of the microbiome included in the model, the ones with the highest importance in the probability forest were selected for further analysis. The loadings of these principal components were clustered using an unsupervised hierarchical clustering algorithm to visualize some of the most influential OTUs’ impact on these principal components.

### Data availability

Raw files for bacterial and fungal amplicons for each sample are available in NCBI under BioProject accession code: PRJNA699261.

## ACKNOWLEDGMENTS

A.A. is a cofounder, and N.I. and D.A. are current employees of Biome Makers, Inc. A.D., J.D., V.T. are current employees of Crop Science Division in Bayer. I.B. was an employee of Biome Makers, Inc. at the time of designing the work, but he is now an independent researcher at the Complutense University of Madrid. The soil applied biological used in this article is commercialized by Bayer CropScience LP under the name Minuet. Some authors (N.I., I.B., D.A., and A.A.) have a US pending patent application in relation to this work: USPTO Serial Number 17/119,972. Some authors (N.I., D.A., and A.A.) have US provisional patent applications in relation to this work: USPTO Serial Numbers 63/143,159; 63/143,534; and 63/143,600. Authors received funding from Bayer CropScience LP for this project. We would like to thank Dr. Lauren Cline and Dr. Punita Juneja from Crop Science Division in Bayer for valuable discussions, and Dr. Francisco Ossandon, Mr. Luis Lopez and Mr. Pablo Ahumada from Biome Makers for help with the bioinformatics processing of raw data.

## AUTHOR CONTRIBUTIONS

I.B., A.D., J.D., V.T., and A.A. conceived and designed the work; N.I. contributed to the development of the analytical pipelines, and built the data and computational infrastructure; N.I., I.B., and D.A. performed and supervised data analysis; N.I., I.B., D.A., and A.A. wrote the manuscript. All authors reviewed and approved the submitted version.

## Supplementary Figures Legends

**Figure S1.** Taxonomic composition of soil samples across locations and sampling times. (A) Most abundant bacterial genera identified. (B) Most abundant fungal genera identified. T0 - before planting; T1 - one month after planting; T2 - two months after planting; T3 – after harvest.

**Figure S2.** Taxonomic assignment and their relationship with yield (fold change values) of the OTUs contributing to the ten most important principal components of the beta-diversity ordination generated for the yield predictive model.

## Supplementary Table Legends

**Table S1.** Full PERMANOVA for bacterial and fungal beta-diversity.

**Table S2.** Wilcoxon rank-sum test for bacterial and fungal alpha-diversity of control vs. treated samples.

**Table S3.** Wilcoxon rank-sum test for bacterial and fungal alpha-diversity of T0 vs. T3 samples.

**Table S4.** Significant differential abundance of bacterial and fungal OTUs of control vs. treated samples at T1 vs. T0 and T2 vs. T0.

**Table S5.** Network property changes of bacterial and fungal communities of control vs. treated samples at T1 vs. T0 and T2 vs. T0.

**Table S6.** Importance of variables in Random Forest yield predictive models.

## References

1. Haverkort AJ, Struik PC. 2015. Yield levels of potato crops: recent achievements and future prospects. Field Crops Res 182:76–85.

2. United Nations. 2019. World Population Prospects 2019, viewed September 30th 2020, <https://population.un.org/wpp/>

3. Van Ittersum MK, Cassman KG, Grassini P, Wolf J, Tittonell P, Hochman Z. 2013. Yield gap analysis with local to global relevance-a review. Field Crops Res 143:4–17.

4. Qin S, Zhang J, Dai H, Wang D, Li D. 2014. Effect of ridge–furrow and plastic-mulching planting patterns on yield formation and water movement of potato in a semi-arid area. Agric Water Manag 131:87–94.

5. Daryanto S, Wang L, Jacinthe, PA. 2016. Drought effects on root and tuber production: A meta-analysis. Agric Water Manag 176:122–131.

6. Zarzecka K, Gugała M, Sikorska A, Grzywacz K, Niewęgłowski M. 2020. Marketable Yield of Potato and Its Quantitative Parameters after Application of Herbicides and Biostimulants. Agriculture 10:49.

7. Oliveira JS, Brown HE, Gash A, Moot DJ. 2016. An explanation of yield differences in three potato cultivars. Agron J 108:1434–1446.

8. Reiter B, Pfeifer U, Schwab H, Sessitsch A. 2002. Response of endophytic bacterial communities in potato plants to infection with *Erwinia carotovora* subsp. Atroseptica. Appl Environ Microbiol 68:2261–2268.

9. Rasche F, Velvis H, Zachow C, Berg G, Van Elsas JD, Sessitsch A. 2006a. Impact of transgenic potatoes expressing anti□bacterial agents on bacterial endophytes is comparable with the effects of plant genotype, soil type and pathogen infection. J Appl Ecol 43:555–566.

10. Rasche F, Hödl V, Poll C, Kandeler E, Gerzabek MH, Van Elsas JD, Sessitsch A. 2006b. Rhizosphere bacteria affected by transgenic potatoes with antibacterial activities compared with the effects of soil, wild-type potatoes, vegetation stage and pathogen exposure. FEMS Microbiol Ecol 56:219–235.

11. Weinert, N, Meincke, R, Gottwald, C, Heuer, H, Gomes, N.C, Schloter, M, Berg, G. and Smalla, K. 2009. Rhizosphere communities of genetically modified zeaxanthin-accumulating potato plants and their parent cultivar differ less than those of different potato cultivars. Appl Environ Microbiol 75:3859–3865.

12. Weinert N, Meincke R, Gottwald C, Heuer H, Schloter M, Berg G, Smalla K. 2010. Bacterial diversity on the surface of potato tubers in soil and the influence of the plant genotype. FEMS Microbiol Ecol 74:114–123.

13. İnceoğlu Ö, Al-Soud WA, Salles JF, Semenov, AV, van Elsas JD. 2011. Comparative analysis of bacterial communities in a potato field as determined by pyrosequencing. PloS One 6:e23321.

14. İnceoğlu Ö, Salles, JF, van Elsas, JD. 2012. Soil and cultivar type shape the bacterial community in the potato rhizosphere. Microb Ecol 63:460–470.

15. Kõiv V, Roosaare M, Vedler E, Kivistik PA, Toppi K, Schryer DW, Remm M, Tenson T, Mäe A. 2015. Microbial population dynamics in response to *Pectobacterium atrosepticum* infection in potato tubers. Sci Rep 5:11606.

16. Jeanne T, Parent SÉ, Hogue R. 2019. Using a soil bacterial species balance index to estimate potato crop productivity. PloS One 14:e0214089.

17. NRCS, Soil Health, NRCS’s portal, Natural Resources Conservation Service of the United States Department of Agriculture, viewed September 30th 2020, <https://www.nrcs.usda.gov/wps/portal/nrcs/main/soils/health/>

18. Tahat M, Alananbeh K, Othman Y, Leskovar D. 2020. Soil Health and Sustainable Agriculture. Sustainability 12:4859.

19. Colla G, Nardi S, Cardarelli M, Ertani A, Lucini L, Canaguier R, Rouphael, Y. 2015. Protein hydrolysates as biostimulants in horticulture. Sci Hortic 196:28–38.

20. Van Oosten MJ, Pepe O, De Pascale S, Silletti S, Maggio, A. 2017. The role of biostimulants and bioeffectors as alleviators of abiotic stress in crop plants. Chem Biol Technol Agric 4:5.

21. Grand View Research. 2018. Biostimulants Market Size, Share & Trends Analysis Report By Active Ingredient (Acid Based, Seaweed Extract, Microbial), By Crop Type (Row Crops & Cereals), By Application (Foliar, Soil), And Segment Forecasts, 2018 – 2025.

22. Rajabi-Hamedani S, Rouphael Y, Colla G, Colantoni A, Cardarelli, M. 2020. Biostimulants as a Tool for Improving Environmental Sustainability of Greenhouse Vegetable Crops. Sustainability 12:5101.

23. Naseby D, Pascual J, Lynch J. 2000. Effect of biocontrol strains of *Trichoderma* on plant growth, *Pythium ultimum* populations, soil microbial communities and soil enzyme activities. J Appl Microbiol 88:161–169.

24. Kumar M, Ashraf S. 2017. Role of *Trichoderma* spp. As a biocontrol agent of fungal plant pathogens. In: V. Kumar, M. Kumar, S. Sharma, R. Prasad (Eds.), Probiotics and Plant Health, Springer, Singapore (2017), pp. 497–506.

25. Nelson LM. 2004. Plant growth promoting rhizobacteria (PGPR): Prospects for new inoculants. Crop Manag 3:1–7.

26. Ribaudo CM, Krumpholz EM, Cassán FD, Bottini R Cantore ML, Curá JA. 2006. *Azospirillum* sp. Promotes Root Hair Development in Tomato Plants through a Mechanism that Involves Ethylene. J Plant Growth Regul 25:175–185.

27. Souza R, Ambrosini A, Passaglia LMP. 2015. Plant growth-promoting bacteria as inoculants in agricultural soils. Genet Mol Biol 38:401–419.

28. Singh VK, Singh AK, Singh PP, Kumar A. 2018. Interaction of plant growth promoting bacteria with tomato under abiotic stress: a review. Agric Ecosyst Environ 267:129–140.

29. Remans R, Croonenborghs A, Gutierrez RT, Michiels J, Vanderleyden J. 2007. Effects of plant growth-promoting rhizobacteria on nodulation of *Phaseolus vulgaris* L. are dependent on plant P nutrition. Eur J Plant Pathol 119:341–351.

30. Rodrigues EP, Rodrigues LS, de Oliveira ALM, Divan-Baldani VL, dos Santos-Teixeira KR, Urquiaga S, Massena Reis, V. 2008. *Azospirillum amazonense* inoculation: effects on growth, yield and N2 fixation of rice (*Oryza sativa* L.). Plant Soil 302:249–261.

31. Arif I, Batool M, Schenk PM. 2020. Plant Microbiome Engineering: Expected Benefits for Improved Crop Growth and Resilience. Trends Biotechnol. In Press.

32. Toju H, Peay KG, Yamamichi M, Narisawa K, Hiruma K, Naito K, Fukuda S, Ushio M, Nakaoka S, Onoda Y, Yoshida K, Schlaeppi K, Bai Y, Sugiura R, Ichihashi Y, Minamisawa K, Kiers ET. 2018. Core microbiomes for sustainable agroecosystems. Nat Plants 4:247–257.

33. Ricci M, Tilbury L, Daridon B, Sukalac K. 2019. General principles to justify plant biostimulant claims. Front Plant Sci 10:494.

34. Cavaglieri l, Orlando J, Etcheverry M. 2009. Rhizosphere microbial community structure at different maize plant growth stages and root locations. Microbiol Res 164:391–399.

35. Wang W, Luo X, Chen Y, Ye X, Wang H, Cao Z, Ran W, Cui Z. 2019. Succession of Composition and Function of Soil Bacterial Communities During Key Rice Growth Stages. Front Microbiol 10:421.

36. Carrino-Kyker SR, Coyle KP, Kluber LA, Burke DJ. 2020. Fungal and Bacterial Communities Exhibit Consistent Responses to Reversal of Soil Acidification and Phosphorus Limitation over Time. Microorganisms 8:1.

37. Qiao J, Yu X, Liang X, Liu Y, Borriss R, Liu Y. 2017. Addition of plant-growth-promoting *Bacillus subtilis* PTS-394 on tomato rhizosphere has no durable impact on composition of root microbiome. BMC Microbiology 17:131.

38. Correa OS, Montecchia MS, Berti MF, Fernández-Ferrari MC, Pucheu NL, Kerber NL, García AF. 2009. *Bacillus amyloliquefaciens* BNM122, a potential microbial biocontrol agent applied on soybean seeds, causes a minor impact on rhizosphere and soil microbial communities. Appl Soil Ecol 41:185–194.

39. Chowdhury SP, Dietel K, Rändler M, Schmid M, Junge H, Borriss R, Hartmann A, Grosch R. 2013. Effects of *Bacillus amyloliquefaciens* FZB42 on Lettuce Growth and Health under Pathogen Pressure and Its Impact on the Rhizosphere Bacterial Community. PLoS ONE 8:e68818.

40. Ortiz-Álvarez R, Ortega-Arranz H, Ontiveros VJ, Ravarani CN, Acedo A, Belda I. 2020. Emergent properties in microbiome networks reveal the anthropogenic disturbance of farming practices in vineyard soil fungal communities. bioRxiv doi:10.1101/2020.03.12.983650.

41. Devictor V, Clavel J, Julliard R, Lavergne S, Mouillot D, Thuiller W, Venail P, Villéger S. Mouquet N. 2010. Defining and measuring ecological specialization. J Appl Ecol 47:15–25.

42. Woo SL, Ruocco M, Vinale F, Nigro M, Marra R, Lombardi N, Pascale A, Lanzuise S, Manganiello G, Lorito M. 2014. *Trichoderma*-based products and their widespread use in agriculture. Open Mycol J. 8:71–126.

43. Van Klompenburg T, Kassahun A, Catal, C. 2020. Crop yield prediction using machine learning: A systematic literature review. Comput Electron Agric 177:105709.

44. U.S. Environmental Protection Agency. 2006. Biopesticide Registration Action Document. Bacillus subtilis Strain QST 713 (PC Code 006479), viewed September 30th 2020, <https://www3.epa.gov/pesticides/chem_search/reg_actions/registration/decision_PC-0064799-Aug-06.pdf>.

45. Pérez-García A, Romero D, De Vicente A. 2011. Plant protection and growth stimulation by microorganisms: biotechnological applications of Bacilli in agriculture. Curr Opin Biotech 22:187–193.

46. Falardeau J, Wise C, Novitsky L, Avis TJ. 2013. Ecological and mechanistic insights into the direct and indirect antimicrobial properties of Bacillus subtilis lipopeptides on plant pathogens. J Chem Ecol 39:869–878.

47. Lundberg DS, Lebeis SL, Paredes SH, Yourstone S, Gehring J, Malfatti S, Tremblay J, Engelbrektson A, Kunin V, Del Rio TG, Edgar RC, Eickhorst T, Ley RE, Hugenholtz P, Tringe SG, Dangl JL. 2012. Defining the core *Arabidopsis thaliana* root microbiome. Nature 488:86–90.

48. Acedo Becares A, Ferrero Fernandez A, Biome Makers Inc. 2018. Microbiome based identification, monitoring and enhancement of fermentation processes and products. U.S. Patent Application 15/779,531.

49. Rognes T, Flouri T, Nichols B, Quince C, Mahé F. 2016. VSEARCH: a versatile open source tool for metagenomics. PeerJ 4:e2584.

50. Edgar RC, Hass BJ, Clemente JC, Quince C, Knight R. 2011. UCHIME improves sensitivity and speed of chimera detection. Bioinformatics 27: 2194-200.

51. Edgar RC 2016. SINTAX: a simple non-Bayesian taxonomy classifier for 16S and ITS sequences. bioRxiv 074161.

52. Glöckner FO, Yilmaz P, Quast C, Gerken J, Beccati A, Ciuprina A, Bruns G, Yarza P, Peplies J, Westram R, Ludwig W. 2017. 25 years of serving the community with ribosomal RNA gene reference databases and tools. J Biotechnol 261:169–176.

53. Nilsson RH, Larsson KH, Taylor AFS, Bengtsson-Palme J, Jeppesen TS, Schigel D, Kennedy P, Picard K, Glöckner FO, Tedersoo L, Saar I. 2019. The UNITE database for molecular identification of fungi: handling dark taxa and parallel taxonomic classifications. Nucleic Acids Res 47:D259–D264.

54. Oksanen J, Blanchet FG, Friendly M, Kindt R, Legendre P, McGlinn D, Minchin PR, O’Hara RB, Simpson GL, Solymos P, Stevens MHH, Szoecs E, Wagner H. 2019. vegan: Community Ecology Package. R package version 2.5-6.

55. Martín-Fernández JA, Hron K, Templ M, Filzmoser P, Palarea-Albaladejo J. 2015. Bayesian-multiplicative treatment of count zeros in compositional data sets. Stat Model 15:134–158.

56. Robinson, MD, McCarthy DJ, Smyth GK. 2010. edgeR: a Bioconductor package for differential expression analysis of digital gene expression data. Bioinformatics 26:139–1401.

57. Veech JA. 2013. A probabilistic model for analysing species co occurrence. Glob Ecol Biogeogr 22:252–260.

58. Csardi G, Nepusz T. 2006. The igraph software package for complex network research. Inter Journal Complex Systems 1695:1–9.

59. Proulx SR, Promislow DEL, Phillips PC. 2005. Network thinking in ecology and evolution. Trends Ecol Evol 20:345–353.

60. Malley JD, Kruppa J, Dasgupta A, Malley KG, Ziegler A. 2012. Probability machines: consistent probability estimation using nonparametric learning machines. Methods Inf Med 51:74–81.

