## Supplemental Figure S1 for "Soil Microbial Composition and Structure Allow Assessment of Biological Product Effectiveness and Crop Yield Prediction"

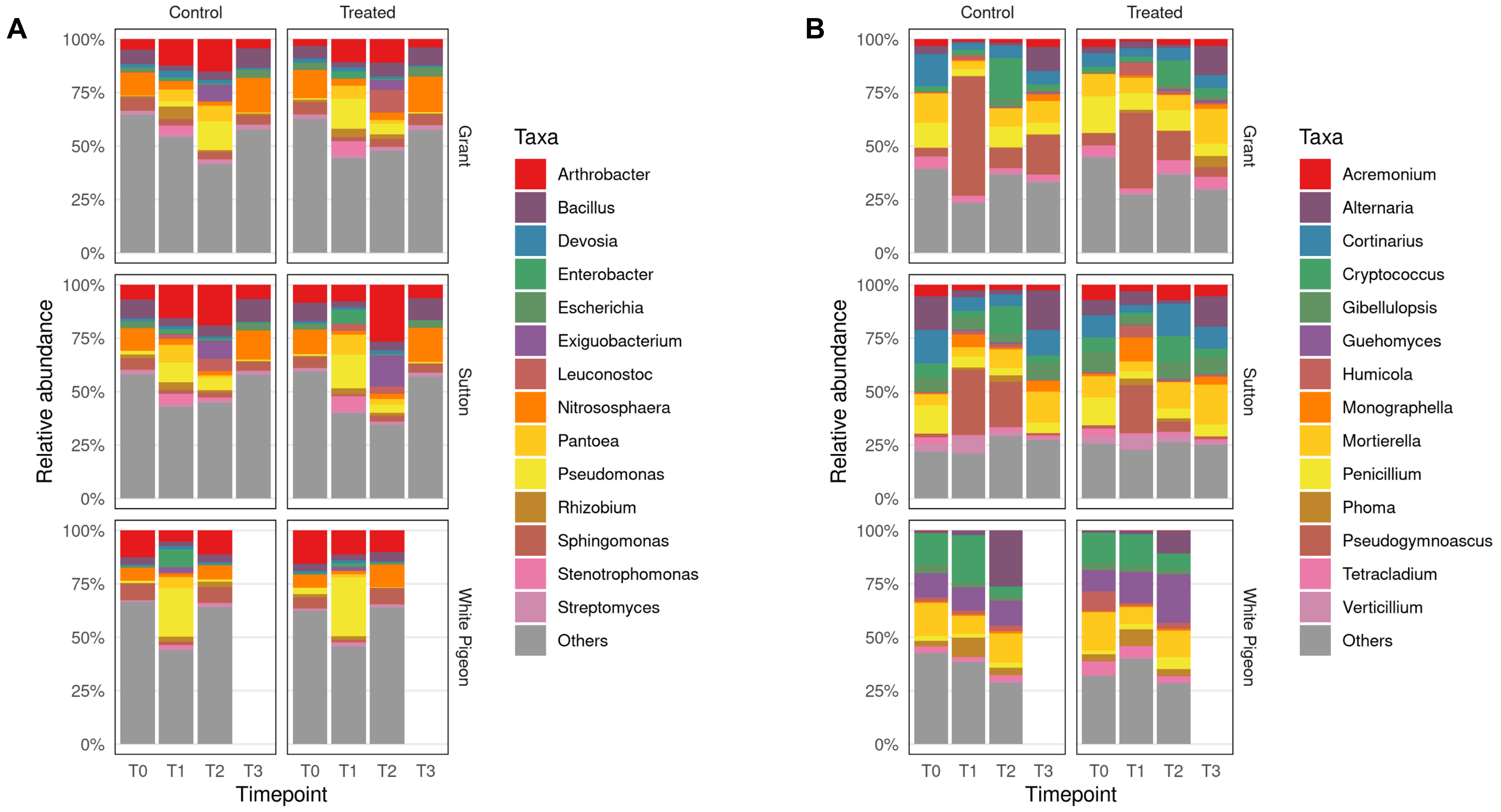


**Figure S1.** Taxonomic composition of soil samples across locations and sampling times. (A) Most abundant bacterial genera identified. (B) Most abundant fungal genera identified. T0 - before planting; T1 - one month after planting; T2 - two months after planting; T3 – after harvest.
