## Supplemental Figure S2 for "Soil Microbial Composition and Structure Allow Assessment of Biological Product Effectiveness and Crop Yield Prediction"

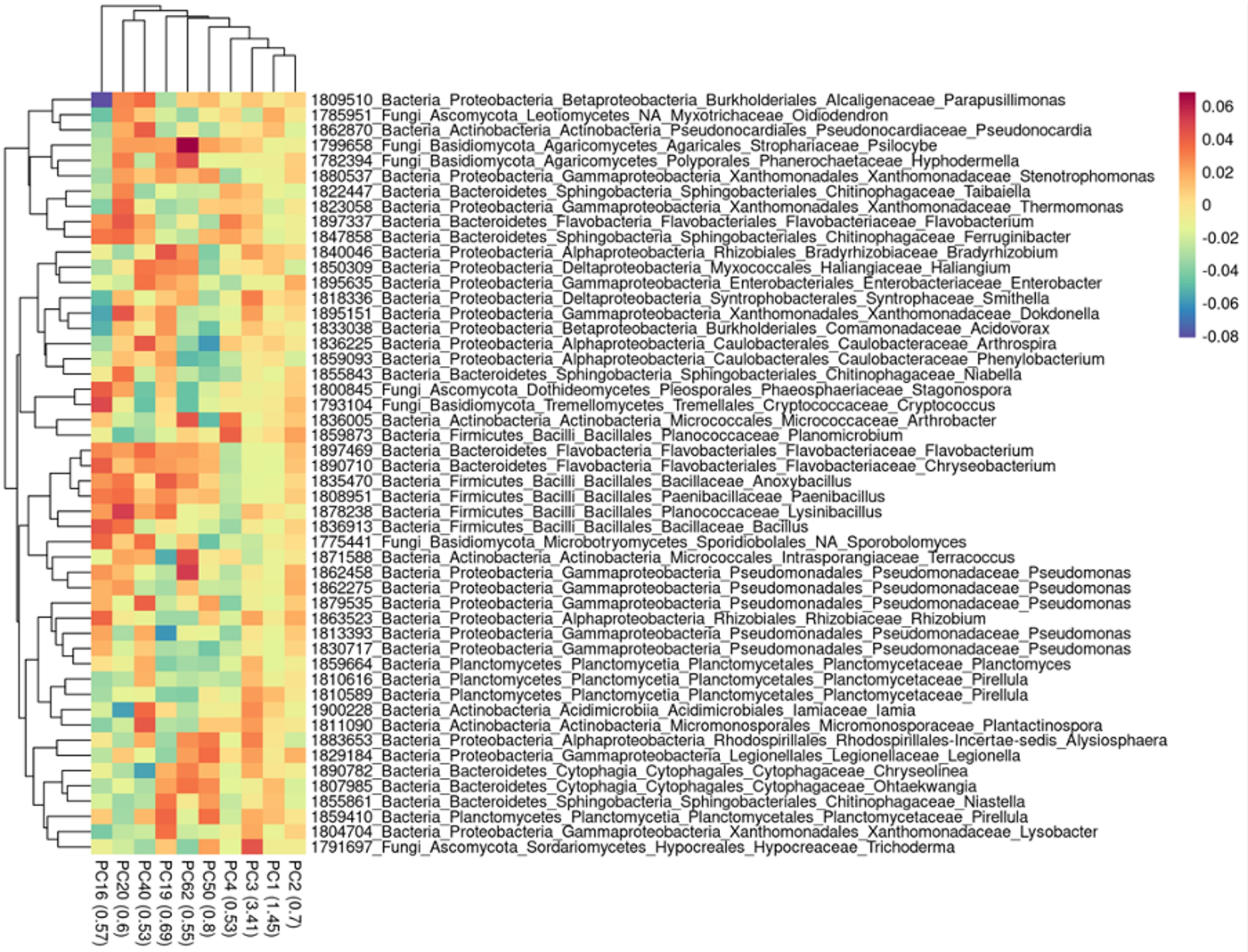


**Figure S2.** Taxonomic assignment and their relationship with yield (fold change values) of the OTUs contributing to the ten most important principal components of the beta-diversity ordination generated for the yield predictive model.
